# Genome sequences of three Konjac mosaic virus (KoMV) variants

**DOI:** 10.1101/2024.02.16.579633

**Authors:** Timo M. Breit, Wim de Leeuw, Marina van Olst, Wim A. Ensink, Selina van Leeuwen, Martijs J. Jonker, Rob J. Dekker

## Abstract

This study explores the genomic diversity of three novel Konjac mosaic virus (KoMV) variants, originating from distinct Zantedeschia (calla lily) commercial cultivars. Virus-derived small RNA sequencing was performed and the complete KoMV variant genome sequences were determined by *de novo* assembly of short reads. All KoMV variants showed substantial nucleotide, as well as protein differences as compared to the KoMV RefSeq sequence.

## Introduction

Potyvirus infection of calla lilies (*Zantedeschia spp*.) can result in various symptoms, such as mosaic and leaf distortion (1, 2). To obtain more insight in the genetic variation of potyviruses in calla lily, we sequenced small RNA from 16 plants of various undisclosed commercial cultivars with symptoms that are indicative for potyvirus infection. Using the siRNA response of the plant, potential infections with known potyviruses were identified (3). We successfully obtained complete genome sequences of three variants of *Konjac mosaic virus* (KoMV; family Potyviridae, genus Potyvirus) from three out of 16 plants. KoMV is a single positive-strand RNA virus with a genome of 9,544 nucleotides and a single open reading frame coding for the characteristic potyviral 350-kDa polyprotein (4).

## Materials and methods

Fresh calla lily leaf material was collected from an anonymous Dutch plant-breeding company situated in the province of North-Holland (The Netherlands) in September 2022. Total RNA was isolated from ±1 cm^2^leaf fragments and enriched for small RNA as described (5). Barcoded small RNA libraries were generated using the Small RNA-Seq Library Prep Kit (Lexogen), according to the supplied instructions. The size distribution of the libraries was assessed using a 2200 TapeStation System with Agilent D1000 ScreenTapes (Agilent Technologies). The libraries were sequenced with 75 cycles on a NextSeq 550 Sequencing System (Illumina) using a NextSeq 500/550 High Output Kit v2.5 (Illumina). First, reads were trimmed using trimmomatic v0.39 (6) [parameters: LEADING:3; TRAILING:3; SLIDINGWINDOW:4:15; MINLEN:19]. Next, full-length KoMV variant genomes were constructed from contigs that were assembled using SPAdes De Novo Assembler (7) with parameter settings: only-assembler mode, coverage depth cutoff 10, and kmer length 17, 18 and 21.

## Results and discussion

We successfully determined the complete genome sequences of three different KoMV variants in three plants with an average depth of coverage of >5,600 and a breadth of coverage of 100% (S07, S13 and S16; Table 1). Each variant exhibits approximately 93.5% sequence similarity when compared to the KoMV RefSeq sequence NC_007913. The single open reading frame of the variants var-UvA-E1-S07/S13/S16 differed 594, 602 and 609 nucleotides from the reference sequence, resulting in 122, 129 and 122 amino acid differences, respectively (Table 1). Among each other, the new KoMV variants showed a sequence identity of 93-99%, with var-UvA-E1-S07 and UvA-E1-S16 resembling each other the most (99%), and UvA-E1-S13 resembling the other variants by 93%. Additionally, when comparing the P1 mature peptide, all three variants had between 9.5 and 11.9% amino acid differences with the reference, whereas the other mature peptides differed only between 0 and 5.8%. This finding supports previous research that suggests the P1 mature peptide is the most divergent of all potyviral proteins (8, 9).

**Table 1.**
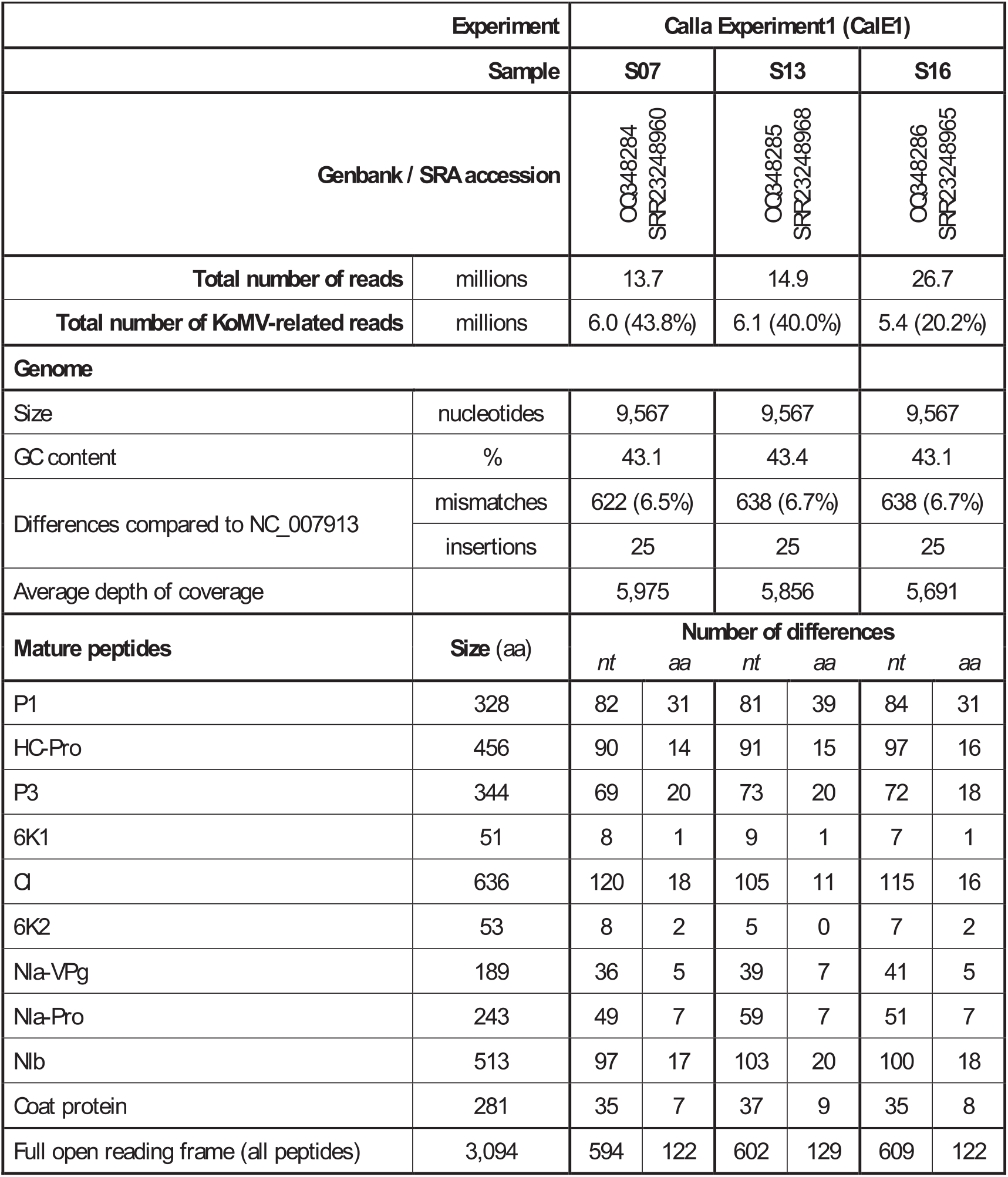
Genome and protein differences of KoMV variants. Nucleotide and amino acid sequences of the mature peptides were obtained from the single open reading frame of each KoMV variant, as annotated in the reference genome NC_007913. The number of differing nucleotides (nt) and amino acids (aa) is shown for each mature peptide compared to the reference genome. New virus genome sequences were constructed from samples with 100% breadth of coverage and high depth of coverage, and these sequences were submitted to GenBank under the GenBank accession numbers indicated. The raw sequence data can be accessed at the NCBI Sequence Read Archive using the SRA accession numbers.

A phylogenetic analysis including all complete KoMV variant genomes from GenBank, along with the three KoMV variants identified in the current study, demonstrates that the KoMV RefSeq NC_007913 forms a distinct clade compared to all other variants (Figure 1). This observation aligns with the deletion of 18 nucleotides at position 3009 in NC_007913, which preserves the reading frame and leads to the removal of the amino acid sequence LSW(L/P/H)GK from the P3 mature peptide.

**Figure 1.**
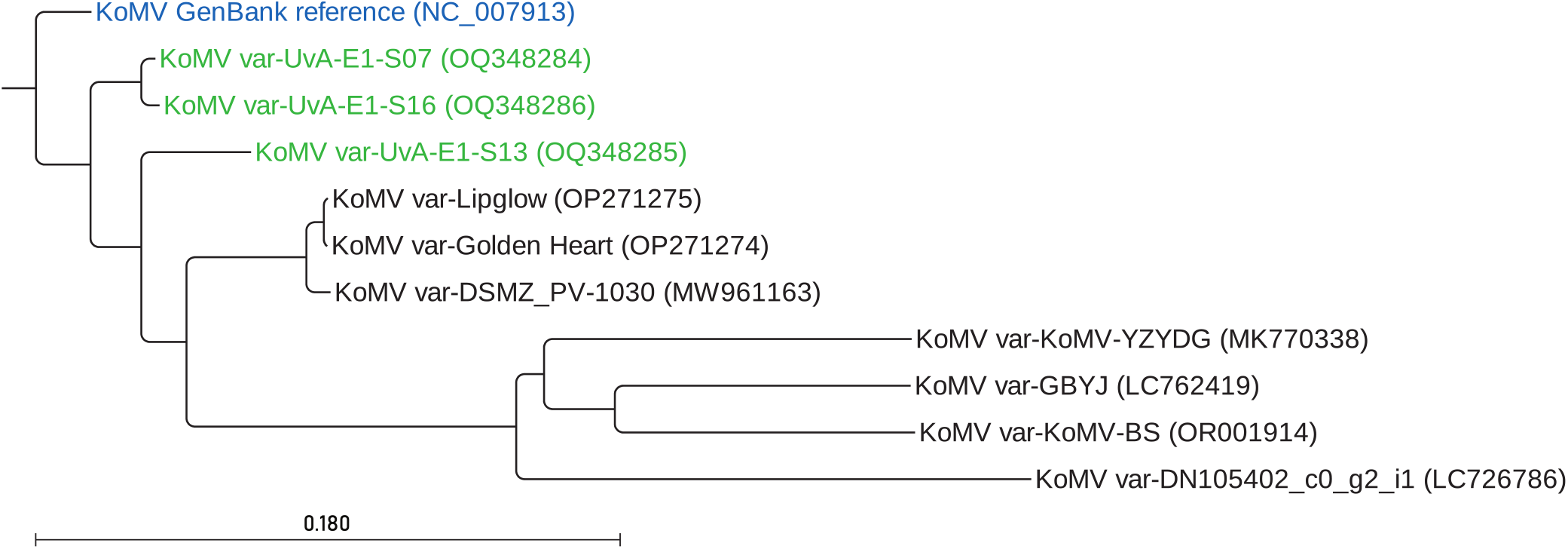
Phylogenetic analysis of KoMV variants. The study included three KoMV variants obtained from the current investigation (represented in green), along with seven KoMV variants sourced from GenBank (represented in black). The KoMV reference genome sequence NC_007913 is shown in blue. The phylogenetic tree was constructed using CLC Genomics Workbench v21 .0.5 software. The tree was generated by initially aligning the genome sequences of the KoMV variants, utilizing the following gap cost settings: open 10, extension 2, and end gap ‘Free’. Subsequently, the Neighbor joining method was employed to construct the tree and Kimura’s two-parameter model (K80) was utilized for estimating genetic distances. To assess the robustness of the tree, bootstrapping was performed with 100 replicates.

## Data availability

The raw sequence reads have been deposited in the NCBI Sequence Read Archive under BioProject accession number PRJNA928251. KoMV variant sequences have been deposited under GenBank accession numbers OQ348284, OQ348285 and OQ348286.

## Acknowledgments

This research was directly and indirectly funded by the Swammerdam Institute for Life Sciences of the University of Amsterdam.

## References

1. Chang YC, Chen YL,Chung FC. 2001. Mosaic disease of calla lily caused by a new potyvirus in Taiwan. Plant Disease 85:1289.

2. Pham K, Langeveld SA, Lemmers MEC, Derks AFLM. 2002. Detection and identification of potyviruses in Zantedeschia. Acta Horticulturae 568:143–148.

3. Kreuze JF, Perez A, Untiveros M, Quispe D, Fuentes S, Barker I, Simon R. 2009. Complete viral genome sequence and discovery of novel viruses by deep sequencing of small RNAs: a generic method for diagnosis, discovery and sequencing of viruses. Virology 388(1):1–7.

4. Nishiguchi M, Yamasaki S, Lu XZ, Shimoyama A, Hanada K, Sonoda S, Shimono M, Sakai J, Mikoshiba Y, Fujisawa I. 2006. Konjak mosaic virus: the complete nucleotide sequence of the genomic RNA and its comparison with other potyviruses. Arch Virol 151(8):1643–50.

5. Breit TM, de Leeuw W, van Olst M, Ensink WA, van Leeuwen S, Jonker MJ, Dekker RJ. 2023. Genome sequences of 10 new carnation mottle virus variants. Microbiol Resour Announc 12(9): e00189–23.

6. Bolger AM, Lohse M, Usadel B. 2014. Trimmomatic: a flexible trimmer for Illumina sequence data. Bioinformatics. 30(15): 2114–2120.

7. Prjibelski A, Antipov D, Meleshko D, Lapidus A, Korobeynikov A. 2020. Using SPAdes De Novo Assembler. Curr Protoc Bioinformatics 70(1):e102.

8. Adams MJ, Antoniw JF, Fauquet CM. 2005 Molecular criteria for genus and species discrimination within the family Potyviridae. Arch Virol 150(3):459–479.

9. Mengual-Chuliá B, Bedhomme S, Lafforgue G, Elena SF, Bravo IG. 2016. Assessing parallel gene histories in viral genomes. BMC Evol Biol 16:32.

